# Genomic footprint of a shared Type 5 prophage in ‘*Candidatus* Liberibacter asiaticus’ and ‘*Candidatus* Liberibacter africanus’, two destructive bacterial pathogens of citrus huanglongbing

**DOI:** 10.1101/2025.05.23.655732

**Authors:** Frédéric Labbé, Claudine Boyer, Fernando Clavijo-Coppens, Blandine Benoist, Patrick Turpin, Santatra Ravelomanantsoa, Olivier Pruvost

## Abstract

Huanglongbing (HLB) is a bacterial disease that affects citrus trees and is considered the most severe citrus disease in the world. The three HLB-associated ‘*Ca.* Liberibacter’ species harbor prophage regions which have been described to play critical roles in their evolution and biology. In this study, we assembled and characterized the accessory genome of the first circular *de novo* ‘*Candidatus* Liberibacter asiaticus’ (CLas) assembly (V1R1) from Réunion, one of the sparse areas worldwide hosting CLas and ‘*Ca.* Liberibacter africanus’ (CLaf). This 1,272 Mb-long whole-genome harbored 1,129 coding sequences and two complete prophages, including a 37,934 bp-long Type 1 prophage, frequently present in CLas genomes, and a 40,501 bp-long undescribed CLas prophage designated as P-V1R1-5. Comparative genomic approaches suggested that P-V1R1-5 have all the genetic elements to produce new viral particles and revealed that it likely belongs to a new CLas Type 5 prophage. While being reported for the first time in a CLas strain, P-V1R1-5-like prophages were previously identified in CLaf genomes, making it the first evidence of shared prophage-like sequence among HLB-associated ‘*Ca*, Liberibacter’ species. Using PCR amplifications targeting P-V1R1-5, we also showed that 85.7% of the tested CLas strains from Réunion and all tested CLaf strains from Madagascar and Réunion harbored a Type 5-like prophage. The identification of CLas Type 5-like prophages not only expanded our knowledge of CLas genomic diversity in Réunion, but also provided new insights into the role of horizontally transferred elements in the evolution of the sympatric HLB-associated bacteria.

**IMPORTANCE:** Huanglongbing (HLB) is the most severe citrus disease worldwide. The disease is associated with three ‘*Candidatus* Liberibacter’ species harboring prophage regions that have been described to play critical roles in their biology. In this study, we assembled and characterized the accessory genome of the first circular *de novo* ‘*Ca.* Liberibacter asiaticus’ (CLas) assembly from Réunion, one of the very few areas in the world where both of CLas and ‘*Ca.* Liberibacter africanus’ (CLaf) coexist. Comparative genomic approaches demonstrated that this genome harbored two complete prophages, including a new CLas Type 5 prophage that was previously identified in CLaf but was reported for the first time in a CLas strain. This first evidence of shared prophage-like sequences among HLB-associated species expands our knowledge of CLas genomic diversity, but also provides new insights into the role of the accessory genome in the evolution of these bacteria.

## INTRODUCTION

Bacteriophages (phages), viruses that infect bacterial, are considered the most abundant biological entity in the biosphere (1). Phages are typically specific to a single bacterial species, although few examples may suggest a broad-host-range including different species within the same genus (2, 3). Phages cannot replicate on their own and they require a bacterial host cell to highjack its machinery for replication through two mechanisms*, i.e.,* the lytic and lysogenic cycles (4). While the lytic lifestyle assembles and ultimately releases new viral particles by the lysis of the host after cell infection, the lysogenic lifestyle, performed by the so-called temperate phages, integrates the phage DNA into the bacterial chromosome and can remain dormant for a period as a prophage allowing its vertical transmission. Prophage integration phenomenon were identified as the major vehicle for horizontal gene transfer contributing significantly to the diversity of the bacterial gene repertoire (5, 6). However, under certain conditions, the prophage manages to hijack the stress signaling pathways, triggering the initiation of the lytic cycle through genome replication, which results in the release of virions into the environment (7, 8). Prophages have been reported in economically important bacterial pathogens of cultivated crops, such as *Xanthomonas citri* (9, 10), the *Ralstonia solanacearum* species complex (11, 12), *Xylella fastidiosa* (13, 14), and ‘*Candidatus* Liberibacter asiaticus’ (CLas) (15, 16). Prophage genes, integrated by temperate phages into different bacterial cells, can play a critical role in intraspecies diversity and bacterial evolution by providing selective advantages, such as virulence traits, resistance to environmental fluctuations, and defense mechanism (17–19).

CLas, is the most prevalent species among the three Gram-negative phloem-restricted α-proteobacteria associated with citrus Huanglongbing (HLB). The disease is considered the most destructive disease of citrus industries (20), *i.e.,* the world’s largest fruit crop in term of production (21). HLB causes alterations of citrus fruit quality and yield, and the gradual death of branches until the tree dies (20). A detailed description of HLB symptoms is available elsewhere (22, 23). The disease has reduced citrus production in Florida by more than 90% since the early 2000s (24). Two other ‘*Ca.* Liberibacter’ species associated with HLB, *i.e.,* ‘*Ca.* Liberibacter africanus’ (CLaf) reported from Africa, the Arabian peninsula and the Mascarene archipelago, and ‘*Ca.* Liberibacter americanus’ (CLam) in Brazil (25, 26). HLB is transmitted by grafting or by two species of phloem-sap feeding psyllids, *i.e.,* the Asian citrus psyllid (ACP, *Diaphorina citri*) and the African citrus psyllid (AfCP, *Trioza erytreae*) (22, 27). None of these HLB causing species are currently culturable *in vitro* and most discoveries are derived from metagenomic and comparative genomic approaches (28–31). As evidenced by the relatively small size of their genomes compared to other members of the *Rhizobiaceae* family (∼1.26 Mb), these three ‘*Ca.* Liberibacter’ species lack several essential metabolic genes and must import vital compounds from their hosts, such as amino acids and ATP (28, 32). However, despite this significant genome reduction, prophages are the most variable horizontally transferred genes in HLB-associated ‘*Ca.* Liberibacter’ species and can represent up to ∼11% of their genomes (33, 34).

The CLas genomes harbored three complete double-stranded DNA *Podoviridae* phages, *i.e.,* Type 1 (SC1-like or P-YN-1-like), Type 2 (SC2-like or P-GD-2-like), and Type 3 (P-JXGC-3-like) (15, 16, 34–36). Two tandemly arranged prophages (SP1 and SP2) were also reported in a CLam genome (strain São Paulo), and one or two tandemly arranged prophages were reported in CLaf genomes (P-PTSAPSY-1 and P-Zim-1), depending on the studies (34, 37–40). Based on the shared gene components, Tan *et al.,* 2021 were able to classify all these CLas and CLaf prophages into the major *Liberibacter* prophage Type SC (33, 34). CLas genomes also harbored remnant prophages, *i.e.,* Type 4, which are also present in CLaf and CLam and, based on the gene components that are shared, were classify by Tan *et al.,* 2021 into the major *Liberibacter* prophage Type LC2 (33, 41). The highly dynamic components of CLas prophages were used to develop prophage typing systems and differentiate strains for population structure and molecular epidemiology studies around the world, *e.g.,* the United States (42–44), China (45–49), India (50), Brazil (51), and several outermost regions of the European Union (52). For example, CLas prophage types were associated with two different regions in mainland China, *i.e.,* while Type 2 was mainly found in low-altitude regions, Type 1 alone or the combination of Types 1 + 3 were more abundant in high-altitude regions (45, 48). More CLas prophages are likely to be discovered due to the recent developments of next generation sequencing technologies, metagenomics analysis, and comparative genomics approaches.

Interestingly, while prophages are not required for CLas pathogenicity (49, 53), transcriptomic and pathogenicity analyses on CLas prophages carrying strains suggested that phages integrated sequences influence CLas pathogenicity and adaptability to its host plants and insect vectors (35, 36, 54–56). The Type 1 prophages encode lysis genes, *i.e.,* a holin (SC1_gp110) and an endolysin (SC1_gp035), and may be present *in planta* as a phage particles (15, 35, 57). Pathogenicity analysis and global gene expression profiling of a strain carrying a Type 1 prophage suggested that this lytic activity may participate in limiting the propagation of this strain (36). The Type 2 prophages lacked lytic genes but encode secreted effectors such as a ROS-scavenging peroxidase (SC2_gp095), which inhibited the reactive oxygen-mediated plant defenses induced by CLas infections (15, 56). Global transcriptomic analysis on a CLas strain carrying a Type 2 prophage confirmed the activation of genes involved in the lysogenic conversion of this temperate phage which could reside as a prophage form within the CLas genome (36). The Type 3 prophages carry a restriction-modification (R-M) system that was speculated to provide a defense against Type 1 phage infection (15, 16, 35, 56). Interestingly, according to Y. Zheng et al., (2025), the CLaf prophage P-Zim-1 harbored 46 coding sequences (CDSs), of which 17 CDSs are homologous to the CDSs encoded by Type 1 and Type 2 CLas prophages (58). One of these homologs encodes a secreted effector (V9J15_02130; homolog of SC1_gp095 and SC2_gp095), suggesting that both CLaf and CLas prophages may use similar strategies to counteract the host immune response and contribute to the infection (56). The CLaf prophages P-Zim-1 and P-PTSAPSY-1, which were also suggested to both carry a potential novel CRISPR/Cas system, seemed to contribute to the bacteria defense mechanism against phage infection and plasmid transfer (58).

Surprisingly, despite the high CLas genetic diversity and high variation of symptoms on HLB-affected trees in Réunion, only the Type 1 prophage, together with the remnant Type 4 prophage, were identified in CLas strains from the island (52, 59). In this outermost region of the European Union (EU), HLB was first reported in the 1968 where both CLas and CLaf were subsequently identified in the lowlands and highlands, respectively. Interestingly, at an intermediate altitudinal range (800-950 meter above sea level; masl) both pathogens co-occurred and can sometimes co-infect a single host (60). Despite a massive control program in the 1990s which decreased disease prevalence at very low levels, HLB reemergence was evidenced in 2015 in most of the citrus cultivation regions of the island, but likely occurred earlier (22, 52, 61, 62). A recent study highlighted that CLas currently outperforms CLaf in Réunion (52). ACP and AfCP insect vectors were also found in Réunion and HLB-transmissibility assays revealed that both vector species transmit both bacterial species (22, 27, 63). Réunion island is one of the very few areas in the world where both of CLas and CLaf and their vectors coexist, making this island a place of choice to investigate horizontally transferred elements, interspecific genomic rearrangement dynamics, and adaptation to host plants and insect vectors.

This study aimed at investigating the integrated phage-like sequences present in CLas genomes by assembling and exploring its first circular *de novo* genome from Réunion (V1R1) produced from short- and long-read co-assembly. Two prophages were identified into this 1,271,573 bp single contig *de novo* assembly, including one prophage that belonged to a new Type 5 CLas prophage (P-V1R1-5) which is highly similar to prophages previously identified in CLaf genomes. Our results suggested that a Type 5-like CLas prophages is present in most CLas strains from Réunion and all CLaf strains from Madagascar and Réunion. Genome annotations suggested that P-V1R1-5 is still capable of both lysogenic and lytic cycles. The identification of this first complete interspecies prophage-sequence not only expanded our knowledge of CLas genomic diversity in Réunion, but also provided new insights into the role of the accessory genome on the diversity, evolution, and biology of the sympatric HLB-causing ‘*Ca.* Liberibacter’ species.

## RESULTS

### Sequencing, Phylogeny and Genome Characteristics

A total of 25,934,866 reads, *i.e.,* 4,518393e^9^ bp, were generated from CLas-infected AfCPs, but only 4,509,586 reads were identified as ‘*Ca*. Liberibacter’ using three whole-genome sequences as references (Table S1). While only 398,842 of those reads (391,422 short and 7,420 long reads) mapped on the CLaf genome PTSAPSY, 2,048,491 reads (2,025,532 short and 22,959 long reads) and 2,009,342 reads (1,986,811 short and 22,531 long reads) respectively mapped on the CLas genomes ReuSP1 and JXGC, confirming that V1R1 genome belongs to a CLas strain. However, 240,890 reads (236,182 short and 4,708 long reads) blasted with both species, *i.e.,* correspond to genomic regions that are highly conserved between the two species. The whole-genome assembly of CLas strain V1R1 comprises 1,271,573 bp in a circular chromosome (GenBank: *accessions will be publicly released upon acceptance of our manuscript for publication*) (Table S1), with the G+C content of 36.58%, and with three rRNA operons, 44 tRNAs, and 1,129 CDSs, including 76 putative Sec-dependent effectors (SDEs), which is consistent with the 86 predicted CLas proteins which were experimentally validated to contain signal peptides (64). The V1R1 genome has a mean depth (*i.e.,* the average number of mapped reads at each base of the genome) of 191X and a coverage (*i.e.,* proportion of the genome covered by mapped reads) of 100%. While most of these CDSs (99.1%) encoded proteins that were shared with at least another ‘*Ca*. Liberibacter’ genome, 10 of them were unique to V1R1 (Table S2), including two putative SDEs, *i.e.,* a collagen triple-helix-repeat (DFOLPVBN_00802) and a lipoprotein (DFOLPVBN_00616). Phylogeny analysis, based on 54 CLas whole-genomes, clustered the two CLas genomes from Réunion, including V1R1, into a highly supported monophyletic group (Fig. S1).

### A Réunion CLas Strain Contains a Prophage Region Homologous to CLaf Strains

We found in V1R1 a 17,016 bp long prophage, categorized as questionable (PHASTEST region’s total score = 80; *i.e.,* small sequence size and no attachment sites), which was located in the 805,000-821,414 genomic region. This incomplete prophage sequence encodes 26 putative CDSs, contains 35.31% of GC, belongs to the remnant Type 4 based on its significant similarity with prophage Ishi-1-a (ANI = 0.999), and was thus designated as P-V1R1-4 (Fig. S2). Two intact phages-like sequences, 40,501 bp and a 37,934 bp long, were also predicted in the V1R1 *de novo* CLas assembly from Réunion located in the 136,646-177,146 and 869,619-907,552 genomic regions and named P-V1R1-1 and P-V1R1-5, respectively, following the nomenclature rule in Z. Zheng *et al.* (2017). The shortest P-V1R1-1 phage sequence showed high similarity with the CLas Type 1 prophage P-YN-1 (ANI = 0.970) (Fig. 1A and Fig. S3). Consistently, the MashTree phylogeny showed that it formed a cluster with P-YN-1, suggesting that this prophage region is highly similar to the CLas Type 1. P-V1R1-1 prophage contained 35 putative CDSs (Table S3), and have an average %GC content of 41.09 which is slightly higher than the average 36.58% GC content of V1R1. With the exception of an exonuclease (DFOLPJBN_00832) and a hypothetical protein (DFOLPJBN_00805), most of the proteins encoded by these CDSs (94.3%) were shared with at least one of the proteins encoded by CLas Type 1 prophages, including the P-ReuSP1-1 prophage previously identified in a CLas strain from Réunion by Lu *et al.,* 2021 (Table S4) (52, 59). Interestingly, the longest P-V1R1-5 prophage showed only low homology with the known CLas prophages (ANI = 0.726 for the closest one, *i.e.,* P-JXGC-3). In contrast, it was found highly similar to the three predicted prophages we re-identified in another ‘*Ca.* Liberibacter’ species, *i.e.,* CLaf (ANI = 0.972, 0.988, and 0.990 for P-PTSAPSY-1, P-Ang37-1, and P-Zim-1, respectively) (Fig. 1A). Similarly, this prophage formed a phylogenic cluster with all these CLaf predicted prophages and shared an identical putative attachment site, *i.e.,* TGGTGCACCCGACAG complementary t-RNA sequence repeat, according to our genome scan. The putative attachment site of this P-V1R1-5 prophage was also found in all tested CLas genomes except one (YNHK-2), including a draft CLas genome from Réunion (ReuSP1; 501,982-501,996) (Table S1). The average depth of P-V1R1-5, *i.e.,* 209X, is close to the average depth of the genome suggesting that it is likely to be a single copy element stably integrated into the genome. Altogether, these results suggested that this CLas complete double-stranded DNA *Podoviridae* phage, which is highly similar to the phages previously identified in CLaf, belongs to a new CLas Type 5 prophage and it was thus designated as P-V1R1-5.

**FIG 1.**
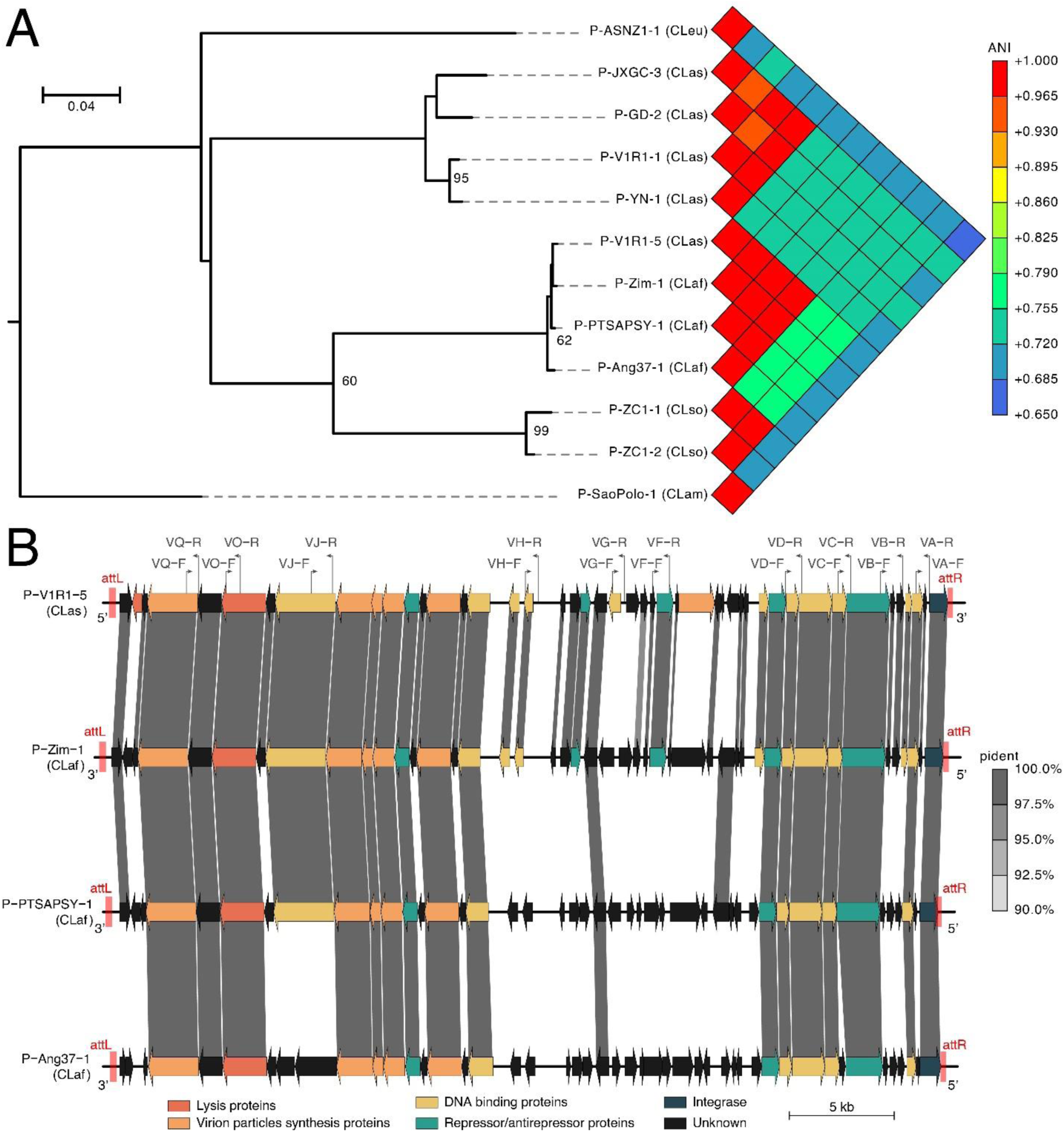
Genomic comparison of the ‘*Candidatus* Liberibacter’ prophages. **(A)** Phylogenetic tree and heatmap of the average nucleotide content (ANI) values. Node support is shown on the phylogenetic tree for bootstrap values greater than 60% (1,000 bootstrap replicate percentage). (**B)** Genomic alignment map of the P-V1R1-5 CLas prophage and the P-Zim-1, P-PTSAPSY-1, and P-Ang37-1 CLaf prophages. The predicted CDSs are indicated by arrows of different colors. The CDSs encoding proteins with a percentage of identity (pident) > 90%, a span > 80%, and e-value < 1e^-3^ are linked by light gray shadings. The locations of the putative attachment sites (*attL* and *attR*) are marked by red regions. The locations of primers specific to the Type 5 prophage P-V1R1-5 are marked by dark-grey lines along with their names and orientations (Table 1). Abbreviations: CLaf: ‘*Ca.* Liberibacter africanus’; CLam: *‘Ca.* Liberibacter americanus’; CLas: *‘Ca.* Liberibacter asiaticus’; CLso: ‘*Ca.* Liberibacter solanacearum’; CLeu: ‘*Ca.* Liberibacter europaeus’.

P-V1R1-5 contained 46 putative CDSs (Fig. 1B and Table S5), and have an average %GC content of 39.73, which is also slightly higher than the average GC content of V1R1. Among its predicted CDSs, 24% are not associated with a known function, while the remaining ones encoded for phage-associated proteins. Based on pairwise comparisons of their amino acid sequences, 40 of the proteins encoded by P-V1R1-5 (87%) were shared with at least one of the proteins encoded by the three described prophages identified in CLaf (Fig. 1B and Table S6), and six were unique to P-V1R1-5 (13%; DFOLPJBN_00126, 00147, 00148, 00153, 00154b, and 00155). However, no proteins encoded by P-V1R1-5 were shared with the proteins encoded by the other prophage Types identified in CLas (Table S7).

Interestingly, according to P-V1R1-5 reannotation using VirClust tool, we identified CDSs encoding repressor/antirepressor proteins involved in prophage dormancy and/or induction, including CDSs encoding RecA (DFOLPJBN_00160 and 00164) and RecX (DFOLPJBN_00136). P-V1R1-5 may also encode CDSs involved in lysogenic conversion, such as a Bro-N family phage antirepressor (DFOLPJBN_00151) and an integrase (DFOLPJBN_00170). However, several CDSs were also involved in the phage lytic cycle, including CDSs encoding lysis proteins, *i.e.,* a holin (DFOLPJBN_00126) and a putative endolysin protein (DFOLPJBN_00130), and virion particles synthesis proteins (DFOLPJBN_00128, 00133:00135, 00138, and 00153). While P-V1R1-5 carried a Cas-like exonuclease (Cas4-domain exonuclease; DFOLPJBN_00163), we did not find any CRISPR nor other Cas-system genes according to the CRISPRCasFinder and DefenseFinder tools and the CasPDB database (*i.e.,* highest percentage of identical positions of 67% for alignment length > 10 amino acids; evalue > 1e^-3^). Other CDSs encoding DNA binding proteins, which are essential for phage DNA synthesis, were also identified in P-V1R1-5 (DFOLPJBN_00132, 00140:00142, 00147, 00159, 00161, 00162, 00167, and 00168). Interestingly, the P-V1R-5 also carried one CDS encoding a putative lipoprotein (DFOLPJBN_00150) which is predicted to contain a typical signal peptide of 25 amino acids long (likelihood = 1.0).

### Distribution and Variants of the P-V1R1-5-Like Prophage Sequences

Based on PCR amplifications screening, most of the tested CLas strains from Réunion (85.7%) contained a P-V1R1-5-like prophage as only four of them (14.3%) did not amplify any of the targeted prophage regions (Fig. 2A and Table S8). Among these strains, we identified five different prophage PCR profiles (profiles 1-5). While the first and second PCR profiles were respectively observed in eight (28.6%) and 13 (46.4%) CLas samples from Réunion, the remaining PCR profiles were only observed once. By contrast, all tested CLaf strains from Réunion and Madagascar amplified at least 8 of the 10 P-V1R1-5 prophage targeted regions, *i.e.,* PCR profiles 2 and 4 were observed in seven (63.6%) and four (36.4%) CLaf strains, respectively (Fig. 2). Interestingly, the PCR profiles reported in both islands always involved an amplification at the six most conserved targeted prophage regions, *i.e.,* the genomic regions amplified by VC, VD, VH, VJ, VO and VQ (Fig. 1B, Fig. 2, and Table 1).

**FIG 2.**
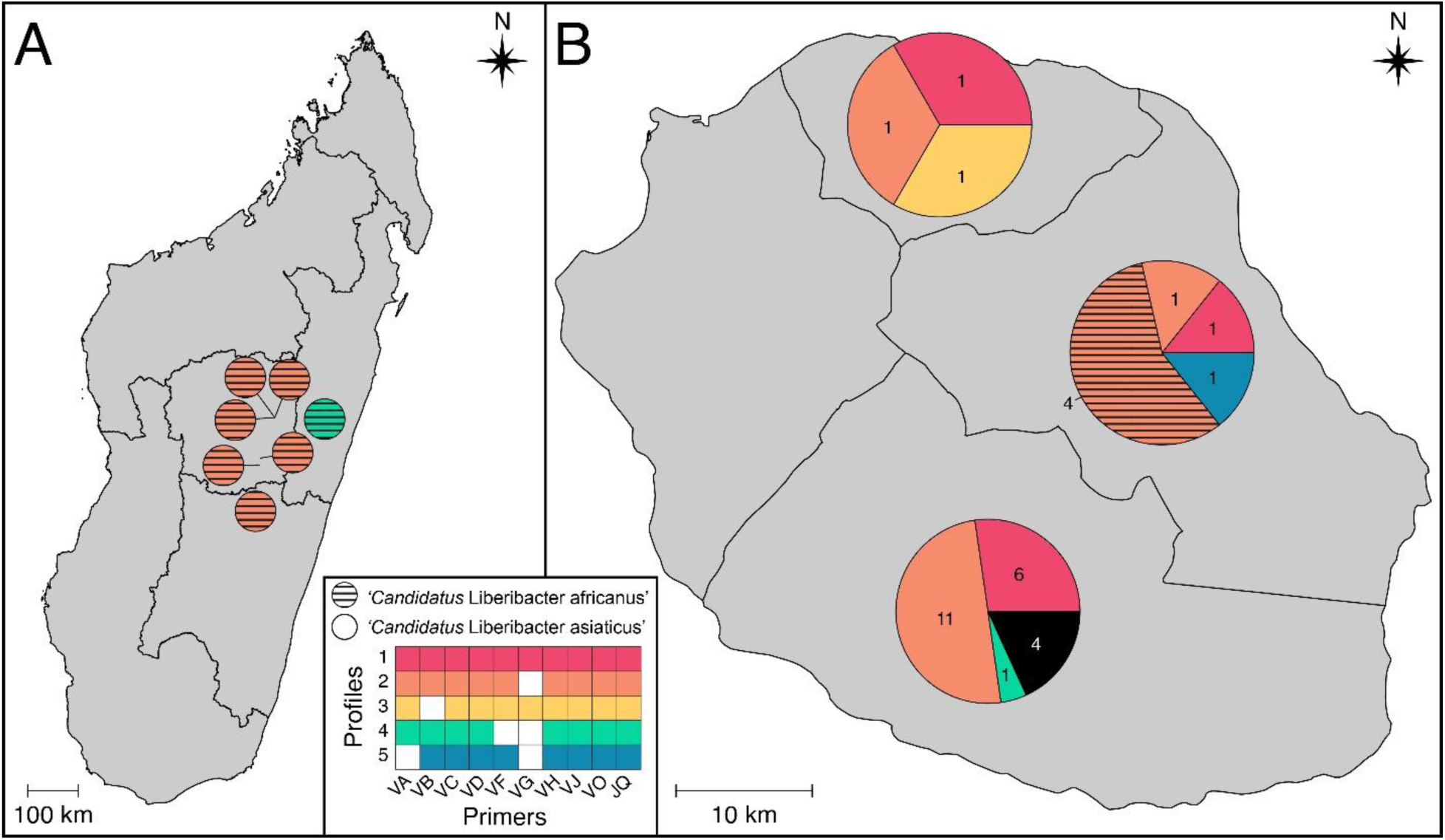
Distribution of the P-V1R1-5-like sequences in Madagascar (A) and Réunion (B). The CLas and CLaf samples harboring different prophage PCR profiles are indicated by different colors. While each circle in Madagascar indicated a PCR profile at a sampling site, each pie chart in Réunion indicated the frequency of the prophage PCR profiles per region. The four CLas samples from Réunion that did not amplify any of the targeted prophage regions are indicated in black. See Fig. 1, Table 1, and Table S7 for more details about the primers. The geographical distribution of the samples was generated using QGIS v.3.22.4 -Białowieża.

**Table 1.**
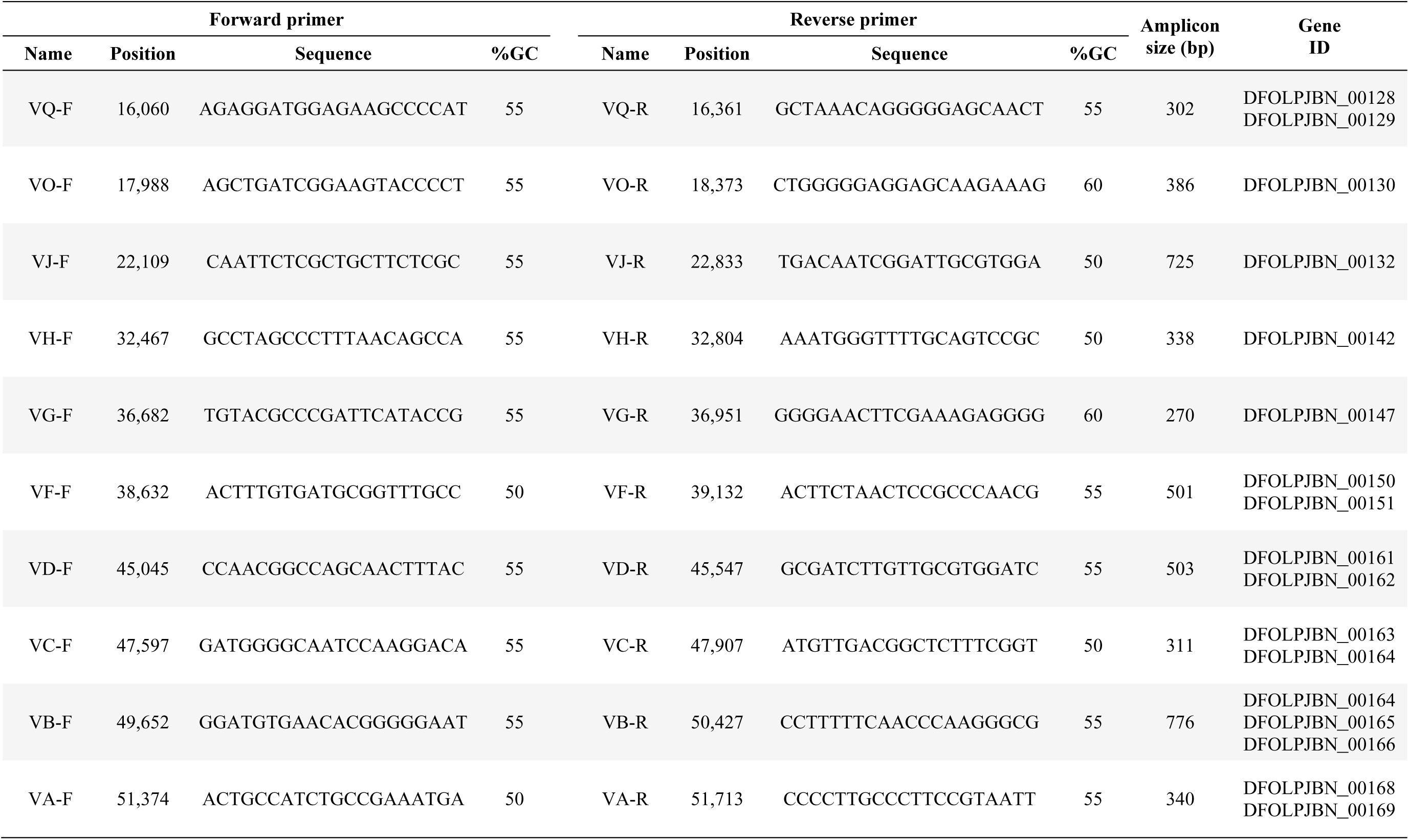
General information of designed primers specific to the ‘*Candidatus* Liberibacter asiaticus’ Type 5 prophage P-V1R1-5 (Fig. 1).

## DISCUSSION

Using *de novo* assembly methods, we assembled the first circular *de novo* whole-genome of a CLas V1R1 strain from Réunion. This whole-genome harbored CLas Type 1 and Type 4 prophages (P-V1R1-1 and P-V1R1-4, respectively), as well as a prophage-like sequence P-V1R1-5 spanning 40,501 bp long and newly described in this study as a new CLas Type 5 prophage. Before this, P-V1R1-5-like prophages had never been reported in other CLas genomes, but homologs have previously been reported in CLaf ones (P-Ang37-1, P-PTSAPSY-1, and P-Zim-1) (38, 58, 66). PCR amplifications screening P-V1R1-5 regions demonstrated that all 11 tested CLaf strains from Réunion and Madagascar, and 24 (85.7%) of the CLas tested strains from Réunion may carried a complete or variants of this prophage. As Pruvost *et al.,* 2024 showed that all 16 CLas strains from Réunion always carried the CLas Type 1 prophage in combination with the remnant Type 4 prophage (52), this new CLas Type 5 integrated phage could be an interesting alternative to differentiate strains in the island using prophage typing systems. Consistently with reported variants for the Types 1, 2 and 3 prophages (37, 42), five Type 5 prophage PCR profiles were identified in the tested strains, suggesting that strains may carry smaller part of this prophage, or that this prophage has evolved enough that most of its genes cannot be amplified by our designed primers. The frequent absence of PCR amplification in the most polymorphic targeted prophage regions gives more weight to the latter hypothesis than to the former, but new genomic resources will be needed to disentangle between these two hypotheses. All prophage regions corresponding to the Type-5 CLas prophage harbored of repeated sequences (TGGTGCACCCGACAG), suggesting the presence of a common putative attachment site. The presence of this Type 5 prophage putative attachment site in near all tested CLas genomes suggests selection for the conservation of integration sites as a means of promoting lysogeny over lysis and facilitating long-term coevolution of temperate phages and bacteria. This heterogeneity of the Type 5 prophage-sequences in the CLaf and CLas strains suggested that both species may share genetic features that allow frequent and independent infection by P-V1R1-5-like active phages. Interestingly, despite the Type 4 and other remnant phages-like sequences were also present in distinct *‘Ca.* Liberibacter’ species (33, 39, 41), the Type 5 prophage provided the first evidence of an interspecies complete prophage-like sequence among HLB-associated ‘*Ca*, Liberibacter’ species.

Despite HLB-associated ‘*Ca.* Liberibacter’ species have evolved towards small genome sizes, the presence of these prophages in their genome suggested that they may contribute to the fitness of these CLas strains. As CLas strains harboring different types of phages (Types 1 or 2) showed different levels of reproduction patterns and higher virulence in plant (36), the biology of the CLas strains harboring this new Type 5 may also be affected. This CLas P-V1R1-5 prophage harbored 46 potential CDSs, most of which are homologous of those found in the predicted prophages of the publicly available CLaf genomes (58). This scarcity of unique CDSs and the presence of shared prophage putative attachment site among these prophages suggested that this phage was able to recognize and infect both species through the same mechanisms. Consistently with evidence of integration of this phage genome within the several bacterial genomes, its gene composition, including the presence of an integrase, an essential enzyme involved in the temperate phage integration process through the catalyzation of unidirectional site-specific recombination between phage and bacterial chromosomal genomes (67), and the presence of a prophage putative attachment site supported that P-V1R1-5 may be a temperate phage capable of excision and lysogenise of closely related bacterial host strains. However, P-V1R1-5 harbored a putative repressor/anti-repressor complex, and RecA-RecX, suggesting the capability to switch from a lysogenic cycle to a lytic cycle.

The ability of P-V1R1-5 to excise, is supported by the genes encoding anti-repressor proteins, including a gene encoding a Bro-N family phage repressor protein which by inactivation of repressors may allow the induction of the phage lytic cycle. Consistently, P-V1R1-5 harbored genes encoding for lysis proteins, including a unique holin and a putative endolysin involved in the bacterial cell walls lysis to release new viral particles (68). Moreover, endolysins, *i.e.,* LasLYS1 and LasLYS2, have been previously reported in other CLas prophages-regions, including one which confers substantial and long-lasting resistance against Huanglongbing and *Xanthomonas citri* pv. *citri* (35, 69, 70). These results suggesting that the prophage genes involved in lysis, such as those found in P-V1R1-5, could be powerful means to control HLB. However, the examination of non-integrated phage particles in the phloem of CLas and CLaf infected periwinkle samples by transmission electron microscopy will be required to provide further evidence of the Type 5 lysogenic to lytic conversion (15, 57, 71). Based on the presence of a signal peptide, the P-V1R1-5 prophage also harbored a CDS encoding a putative SDE lipoprotein (DFOLPJBN_00150). Membrane lipoprotein Ltp encoded by a temperate phage residing in *Streptococcus thermophiles* could prevent superinfection, *i.e.,* infection of the lysogenic host cell by other phage through blocking DNA injection into the host cytoplasm (72, 73). While the absence of homologs for this CDS in other α-proteobacteria cannot provide more information on its function, the presence of this signal peptide in Type 5 prophages suggested that they could be secreted and potentially contribute to the bacterial virulence or protection. However, *Escherichia coli*-based PhoA assay or transcriptomic and pathogenicity analyses will be required to confirm the expression, extracellular secretion, and role of this potential virulence factor carried by P-V1R1-5.

Based on our analyses, while the 46 predicted proteins of P-V1R1-5 were highly similar to the predicted proteins of the known CLaf prophages, none of them were shared with the proteins of the known CLas prophage Types. These results, together with the absence of Type 1 or Type 2 homologous genes encoding peroxidase in P-V1R1-5 and P-Zim-1, do not support previous results describing 17 CDSs of P-Zim-1 which were homologous to Type 1 / Type 2, including CDSs encoding glutathione peroxidase (V9J15_02125, SC1_gp100, SC2_gp100) and peroxidase (V9J15_02130, SC1_gp095, SC2_gp095) (58). The absence of these prophage-encoded peroxidases, which have been reported to be horizontally acquired secreted effector in CLas that suppresses plant defense (56), encourages us to reconsider the fact that both CLaf and CLas prophages may use similar strategies to counteract the host immune response and contribute to the CLas infection. Similarly, no CRISPR, and only one Cas gene, were found in P-V1R1-5 and in the prophages predicted in the publicly available CLaf genomes. These results did not support the identification of a novel CRISPR/Cas system in the Zim and PTSAPSY CLaf genomes (58). A single complete prophage region was predicted in each CLaf and CLam tested publicly available genome. While consistent with the complete prophage region (P-Zim-1) recently reported in the Zim genome (58), these results are not consistent with the two tandemly aligned prophage segments which were previously identified in the PTSAPSY CLaf genome (38) and in the São Paulo CLam genome (SP1 and SP2) (39). However, the length and gene composition, functions and similarities of these single complete predicted prophages highly suggested that these strains harbored a single complete prophage. Altogether, these results highlighted the necessity to use stricter rules when assigning gene functions based on DNA or amino-acid sequence homologies and to use the most recent tools when conducting CRISPR and Cas protein predictions. These results also illustrated the importance of combining the most recent prophage sequence detection, virus genome annotation, and putative attachment site scanning tools when predicting putative complete or functional prophage sequences within bacterial genomes.

Despite sharing similar targets, only sympatric ‘*Ca.* Liberibacter’ host species would be infected by this Type 5 prophage and show evidences of shared phage-like sequences. Indeed, the transfer of genetic material from one bacterium to another via a phage (*i.e.,* transduction) does not require physical contact between the donating and receiving cells (which occurs in conjugation), but it requires that these cells share similar habitats. While CLas is highly studied worldwide, signatures of this Type 5 prophage were only found in the CLas strains from Réunion, *i.e.,* one of the few places in the world where two HLB-associated ‘*Ca*. Liberibacter’ species, CLas and CLaf, and both vectors, ACP and AfCP, are present. The sympatric areas of these species (∼800-950 masl) likely allow relative frequent interactions between these species, as previously illustrated by the CLas/CLaf co-infections of a single host (60), likely explaining this first description of a shared prophage-sequence within the HLB-associated ‘*Ca*. Liberibacter’ species. Several types of interactions could be at the origin of these phage insertion on both bacterial species, including CLas/CLaf coinfections of a single host plant or insect vector where the temperate phage is present as infectious particle (60). In Brazil, despite both CLas and CLam were reported in the São Paulo State and that some trees were infected simultaneously with the two species (74, 75), no evidence of shared prophages were reported between these ‘*Ca*. Liberibacter’ species. However, the predominance of CLas in this region, especially after the 2008–2009 season (76), suggest that interactions between these species may not occur as frequently as those between CLaf and CLas in Réunion. As only a few circular *de novo* genomes were generated for the HLB-associated ‘*Ca*. Liberibacter’ species worldwide, more shared ‘*Ca*. Liberibacter’ prophage-like sequences are likely to be discovered. This includes small single-stranded DNA (ssDNA) *Microviridae* prophages, such as the CLasMV1 which was recently identified in the genomes of CLas strains from China (77), but was not found in V1R1. Overall, our study emphasizes the need of new genomic resources, especially circular *de novo* genomes, to better describe and understand the pangenome of the HLB-associated ‘*Ca*. Liberibacter’ species.

## MATERIALS AND METHODS

### Insect Material

As they have been reported to carry the highest bacterial titers (27), CLas-infected African citrus psyllid (AfCP; *Trioza erytreae*) adults were retrieved from Reynaud *et al.* (2022) HLB transmission assays (Fig. S4). Briefly, AfCP nymphs from HLB-free laboratory colonies were left to feed and develop during 14 days on CLas-infected detached leaves from one field-collected plant (V1R1) of the ForEl 41 cultivar (a hybrid between mandarin cv. Fortune and tangor cv. Ellendale) grown in a specific orchard at 160 masl in Saint-Pierre, Réunion. Newly emerged AfCP adults were transferred on healthy Volkamer lemon (*C. limonia* Osbeck ‘Volkameriana’) detached receptor leaves for 21 days to allow bacterial multiplication and were then preserved at -80°C for further tests.

### DNA Extraction and Sequencing

Total DNA of CLas-infected AfCP was extracted and used for CLas genome sequencing. DNA was extracted from 29 AfCP adults using the Blood and Tissue kit (Qiagen, Courtaboeuf, France) following the manufacturer’s instructions. Upon measurement of sample concentrations with the Qubit dsDNA HS assay kit (Invitrogen, Courtaboeuf, France) on a Qubit fluorometer (Invitrogen), the CLas-infected status of each insect was checked with the real-time PCR assay developed by Li et al., (2006) and by using the GoTaq® qPCR master mix as recommended by the manufacturer (Promega, Charbonnières-les-Bains, France), the StepOnePlus cycler and the Design and analysis software v.2.5 (Applied Biosystems, Courtaboeuf, France). DNA of AfCP adults positive for CLas (Ct ≤ 30) was pooled and 200 ng DNA was used for library preparation. The prepared library was spotted onto eight Flongle Cells R9.4.1 and sequenced with a MinION Mk1B instrument (Oxford Nanopore Technologies, Oxford, UK). Sequencing was run for 24h or until all pores of the Flongle Cell were depleted. Next generation sequencing library preparations and Illumina short-reads sequencing was performed by Genewiz-Azenta Laboratory sequencing platform (Leipzig, Germany) on the Illumina® NovaSeqTM platform (conditions 2 × 150 bp configuration and 10 million read depth).

### Quality Control, Assembly and Annotation

Statistic summary including quality control of a long read sequencing dataset was performed using NanoStat v.1.6.0 (79). We checked short Illumina read data for quality using fastQC v.0.11.8 (80). Nanopore and Illumina adapters trimming was performed with Porechop v.0.2.4 with default settings (https://github.com/rrwick/Porechop) and with trimmomatic v.0.39 (ILLUMINACLIP:NexteraPE-PE.fa:2:30:10 SLIDINGWINDOW:4:15 MINLEN:150) (81), respectively. Trimmed reads were blasted against three whole-genomes using makeblastdb and blastn v.2.12.0+ (82), *i.e.,* a CLas strain from China (GXPSY; Lin et al., 2013), a CLas strain from La Réunion (ReuSP1; Lu et al., 2021), and a CLaf strain from South Africa (PTSAPSY; Lin et al., 2015) (Table S1). Short and long-reads were co-assembled using Unicycler v.0.5.0 (84). We determined the read depth and coverage of the assembled genome using BEDtools v.2.30.0 (85). The assembled genome was annotated using the MicroScope platform and the rapid prokaryotic genome annotation (PROKKA) v.1.14.6 with the protein FASTA file from the MicroScope annotation used for the --proteins option (86, 87). The prediction of the signal peptides, *i.e.,* short amino acid sequences that control protein secretion and translocation, was performed using SignalP v.6.0 (88) to predict the SDEs (64).

### Phylogenetic Relationships

V1R1 *de novo* genome was compared to other CLas genomes obtained from the NCBI assembly database and the figshare repository (https://doi.org/10.6084/m9.figshare.c.5810090.v1). Genomes were aligned with the reference CLas strain GXPSY genome (CP004005) using Minimap2 v.2.24 (89, 90). The single nucleotide variations (SNVs) were called using BCFtools v.1.13 with the haploid model (91). Genomes with a proportion of missing genotypes higher than 30% were discarded using VCFtools v.0.1.16. (92). Variants having a proportion of missing genotypes higher than 20% and a minor allele frequency greater than or equal to 1% were discarded using VCFtools. The remaining filtered dataset included a total of 53 genomes from 10 countries, including two from La Réunion and 30 from the United States, and contained a total of 2,156 bi-allelic SNPs (Table S1). Geneious v.10.2.6 (https://www.geneious.com) were used for genome alignment and visualization (93, 94). Consensus FASTA sequences were created using vcf2fasta_consensus.py (https://github.com/stsmall/An_funestus/tree/master/vcf/). We determined the best nucleotide substitution model using ModelTest-NG v.0.2.0 (95). The phylogenetic relationships among genomes were reconstructed using the maximum likelihood approach implemented in RAxML-NG v.1.1.0 with a TVM+I+G4 substitution model and the node’s support was assessed using 1,000 bootstrap replicates (96). The resulting phylogenetic tree was visualized using FigTree v.1.4.4 (97). We detected the recombinant sequences within the core genome alignment using ClonalFrameML v.1.12 with the ML tree produced by RAxML-NG as the starting tree (98). We discarded the detected recombinant events from the SNP matrix using a custom python script (*ExclRecPos.py*) and VCFtools. However, as none of the bi-allelic SNPs was located in the detected recombinant events, only SNPs due to mutations were used to reconstructed the phylogenetic relationships. All custom python scripts are available at GitHub, https://github.com/fredericlabbe/CLas_Phylogenomics.

### Prophage Characterization

The prophage regions V1R1 were predicted using the PHAge Search Tool with Enhanced Sequence Translation (PHASTEST) (region’s total score ≥ 90) (99–101). We screened for overlaps between the coordinates of the predicted prophage regions and the genome annotation using BEDtools and we also reannotated these predicted prophage regions using VirClust (102). We searched the clustered regularly interspaced short palindromic repeats (CRISPR) and their associated (Cas) proteins within the predicted prophages using the CRISPRCasFinder v.4.2.20 with default parameters (103). We also searched for any known antiviral systems in the predicted prophages using DefenseFinder v.2.0.0 (104), and predicted the Cas protein by blasting the phage predicted proteins against the CasPDB database (http://i.uestc.edu.cn/CASPDB/) (105). When a tRNA was found within a predicted prophage, its putative attachment sites were scanned using a custom python script (*ScanAtt.py*). Pairwise average nucleotide identity (ANI) analysis based on the BLAST algorithm (ANIb) was conducted between the V1R1 predicted prophages and the CLas Type 4 remnant prophage (from the CLas genome Ishi-1 obtained from the NCBI assembly database; Table S1) (41), and 10 other ‘*Ca.* Liberibacter’ prophages. Among these ten ‘*Ca.* Liberibacter’ prophages, three CLas prophages were obtained from the NCBI nucleotide database, *i.e.,* P-YN-1 (also known as SC1), P-GD-2 (also known as SC2), and P-JXGC-3 which were used as representative of Type 1, Type 2, and Type 3, respectively. The remaining seven prophages were predicted using PHASTEST (region’s total score ≥ 90) on the genomes, obtained from the NCBI assembly database, of two other ‘*Ca.* Liberibacter’ species associated with HLB, *i.e.,* CLaf (PTSAPSY, Ang37, and Zim) and CLam (Sao Paulo), and the genomes of two other members of the genus ‘*Ca.* Liberibacter’, *i.e.,* ‘*Ca.* Liberibacter solanacearum’ (CLso; CLso-ZC1) and ‘*Ca.* Liberibacter europaeus’ (CLeu; ASNZ1) (Table S1). A phylogenetic analysis of these prophage regions was also conducted using Mashtree v.1.4.6 (106). The tree was rooted using the identified prophage of the CLam strain Sao Paulo. All V1R1 prophage CDSs were compared to CLas and CLaf prophage CDSs using blastp. Amino acid sequences were considered to share significant similarity when blast results showed a percentage of identical positions > 90%, span > 80%, and an e-value < 1e^-3^. Genomic alignment maps of all prophage sequence regions predicted on V1R1, CLas and CLaf genomes were generated using blastn comparisons and GenoplotR v.0.8.11 (107).

### Prophage-Sequences Diversity and Geographical Distribution

Distribution and composition of predicted phages-like sequences presents on the *de novo* genome assembly V1R1 were explored on HLB-associated ‘*Ca.* Liberibacter’ species presents in Réunion and Madagascar using PCR amplifications screening. Ten pair of primers, designed with Primer3 in Geneious, were selected for PCR amplification specifically targeting distinct regions from this phage (Table 1 and Fig. 1). As controls, we used V1R1-infected or HLB-free Volkamer lemons and AfCPs. Purified genomic DNAs (gDNA) from HLB-infected plants (real-time PCR Cq < 30) were used for prophage identification by PCR amplifications in 28 CLas infected samples from Réunion, four CLaf infected samples from Réunion, and seven CLaf infected samples from Madagascar. While gDNA samples from Réunion were retrieved from Pruvost *et al.,* (2024), gDNA samples from Madagascar were purified from single symptomatic leaves collected between 2019 and 2020 for which the HLB status was determined according to the French official diagnostic protocol (ANSES/LSV/MA 063 – Version 2 – October 2021).

Phage-regions amplicons were assayed at least twice by PCR using P-V1R1-5 specific primers (Table 1). In cases of unclear PCR outcomes, a third replicate was performed. PCR amplifications were performed using the GoTaq® G2 Flexi DNA Polymerase kit (Promega©, Europe) and was made in a final volume of 25 µL composed of: 15.05 µL of water, 5 µL of 5X Colorless GoTaq® Flexi buffer, 1.5 µL of MgCl_2_ solution (25 mM), 0.25 µL of PCR nucleotide mix (10 mM), 0.2 µL of GoTaq® G2 Flexi DNA polymerase solution (5 U/µL), 1 µL of forward primer solution (10 µM), 1 µL of reverse primer solution (10 µM), and 1 µL of DNA (10 ng/µl). PCR amplifications were performed in a Veriti thermocycler (Applied Biosystems, Courtaboeuf, France) under the following conditions: 95°C for 3 min (HotStart activation and DNA initial denaturation), 40 cycles with DNA denaturation at 95°C for 45 s, hybridization of primers at 60°C for 45 s, extension at 72°C for 1 min, final extension at 72°C for 5 min. For each PCR, we used V1R1 and DNase-free water as the positive and negative control, respectively. PCR products were analyzed using QIAxcel advanced instrument (Qiagen), with a 15 pb-3 kb size marker and a 100 pb-2.5 kb alignment marker, and the QIAxel ScreenGel Software v.1.4.0.

## ACKNOWLEDGMENTS

This work was funded by European Union: European regional development fund (ERDF contract GURDT I2016-1731-0006632), DPP SantéBiodiv project, and Interreg V (project EPIBIO-OI); by the Conseil Régional de La Réunion; and by the Centre de coopération Internationale en Recherche Agronomique pour le Développement (CIRAD). Computational work was realized with the support of the ISDM-MESO Platform at the University of Montpellier funded under the CPER by the French Government, the Occitanie Region, the Metropole of Montpellier and the University of Montpellier. We thank Bernard Reynaud and Hélène Delatte for providing us with the CLas-infected AfCP and the CLaf-infected samples from Madagascar, respectively. Laboratory work was conducted on the Plant Protection Platform (3P, IBISA).

## DATA AVAILABILITY

The completed circular genome sequence of ‘*Ca*. Liberibacter asiaticus’ strain V1R1 has been deposited in the GenBank database [*accession will be publicly released upon acceptance of our manuscript for publication*]. The version described in this paper is the first version. The new prophages P-V1R1-1 and P-V1R1-5 have been deposited in GenBank [*accessions will be publicly released upon acceptance of our manuscript for publication*]. All custom python scripts are available at GitHub, https://github.com/fredericlabbe/CLas_Phylogenomics.

